# New algorithm for pearl millet modelling in APSIM allowing a mechanistic simulation of tillers

**DOI:** 10.1101/2023.02.12.528159

**Authors:** Vincent Garin, Erik Van Oosterom, Greg McLean, Graeme Hammer, Tharanya Murugesan, Sivasakthi Kaliamoorthy, Madina Diancumba, Amir Hajjarpoor, Jana Kholova

## Abstract

We present a new algorithm for pearl millet simulation in APSIM. Compared to the actual released model, this new model increases the ability to simulate dynamic tillers by integrating recent progresses about biological understanding of the tillering mechanism. The new algorithm also offers the possibility to have an increased genetic control over key functions like canopy development and tillering through additional genotype related parameters. Next to model description, we also present the parametrization of 9444 and HHB 67-2, two genotypes broadly used in India. Overall, we could show that the new algorithm is able to reconstruct the main plant function like biomass accumulation and tillering. Some margin of improvement remains concerning the simulation of tiller cessation.

## Introduction

Pearl millet is an important crop cultivated on about 30 millions hectares (ha) in more than 30 countries (Jukanti et al., 2016). In India, pearl millet is mainly cultivated during the rainy (kharif) season. Pearl millet is cultivated in the north of Radjastan (A1 zone), the largest production area characterized by a very low level of precipitation (<400 mm/year), the neighbouring regions (south Radjastan, Gujarat, Haryana; A zone) as well as the centre west part of the country (Karnathaka and Maharashtra; B zone; Figure 1). Pearl millet is an important staple food crop for the most vulnerable population of those regions. Pearl millet is resistant to the most adverse agro-climatic conditions where other crops fail to produce economic yield (Yadav et al., 2012). It uses water efficiently and it is tolerant to heat stress. It is a short cycle plant that can also adapt to favourable environments (Yadav and Rai, 2013).

**Figure 1.**
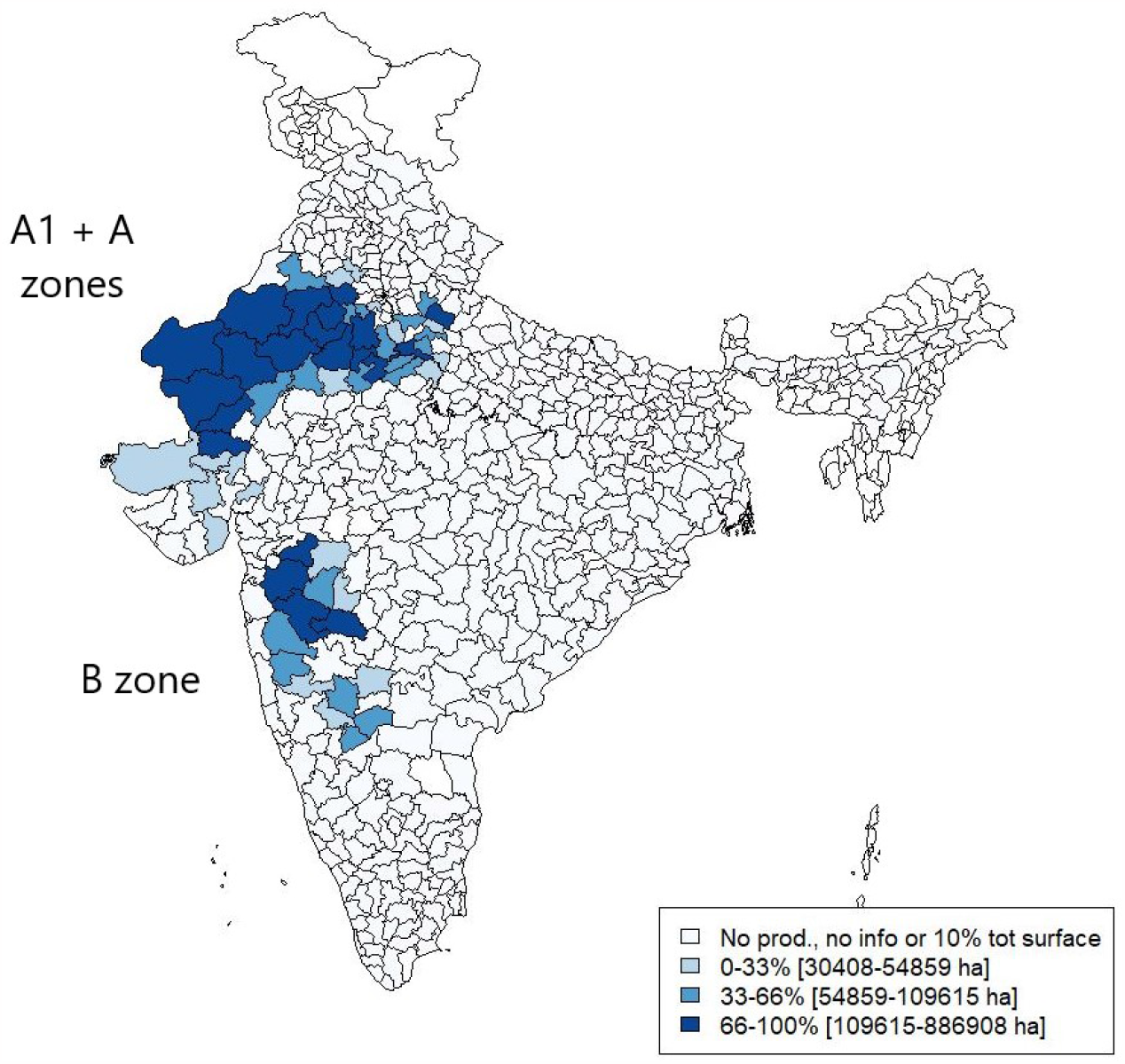
Districts representing 90 % of the total pearl millet cultivated surface area over the period 2000 - 2015

Crop models reconstruct the dynamics of soil-plant-atmosphere continuum of a given agri-system to suit various applications like system characterization and effective system design (Kholova et al., 2020). Crop models can play an important role in supporting the targeted improvement of pearl millet cultivation systems to current and future climate scenarios. This should ultimately increase its socio-economic benefits, which align with several of the sustainable development goals (SDGs). Therefore, the continuous improvement of pearl millet crop models is an important objective.

Unfortunately, compared to other crops like maize, pearl millet crop modelling tends to be an under-researched area. For example, in the agricultural production system simulator (APSIM) environment (Holzworth et al., 2014), since 2001 and the release of a first pearl millet model (Van Oosterom et al., 2001a,b, 2002), few updates have been brought to this initial version. Similarly, the default CERES-Millet model (Godwin et al., 1983) implemented in the decision support system for agrotechnology transfer (DSSAT) environment (Jones et al., 2003) was only recently modified to take better into consideration climatic constraints like drought and heat stress (Singh et al., 2017). Pearl millet crop modelling research was also realized using the SARRA-H model, but this research also starts to date (Sultan et al., 2013). It is therefore important to continue the development of pearl millet model by integrating the latest agronomic and biological discoveries into this valuable corpus of knowledge.

APSIM is a unique modelling environment combining many layers of the agricultural system (soil, weather, plant, management practices) which allows the exploration of the genotype-by-environment-by-management (GxExM) space (Holzworth et al., 2014). In 2001, a first pearl millet model was released in APSIM (Van Oosterom et al., 2001a,b). Taking into consideration pearl millet tillering as an important adaptive mechanism, the authors allowed the model to simulate several tillers by considering the whole plant as an intercrop of the different axes (Van Oosterom et al., 2001b). The need to simulate each tiller individually required a specific function to model the individual leaf area (ILA). The main objective of this work is to describe and evaluate a new algorithm for pearl millet modelling in APSIM. The presented new algorithm builds on the ILA approach but offer more plant specific control over the parameters. In terms of tillering mechanism, the new algorithm proposed by Alam et al. (2014, 2017) is based on available carbohydrate allocation (Kim et al., 2010a,b). Compared to the previous algorithm, the number of tillers does not need to be predetermined by the user and it is not limited. The new algorithm also allows the user to specify plant type specific parameters for the average propensity to tiller and the propensity to tiller given the carbohydrate supply/demand (S/D) ratio. Those modifications makes the tillering mechanism closer to the biological reality, which will have more prediction power over the tiller mechanism and the other emergent properties of the system. The new algorithm also integrates an alternative approach to model the total plant leaf area (TPLA) at the whole plant level. Modelling pearl millet leaf area using TPLA will allow more direct in-silico comparison between pearl millet and competitors like maize for which leaf area is simulated using similar principle.

To evaluate this new algorithm, we parametrized it for two genotypes: 9444 and HHB 67-2, two Indian genotypes representative for intensified (9444) and low-input (HHB 67-2) agriculture. We evaluated the new model prediction ability using data from three experiments and compared the TPLA and ILA approaches. In the next sections we will: a) describe the new algorithm, b) present the experiment data, c) describe the approach used for genotype parametrization and model evaluation, and d) present and discuss the obtained results.

## Material and Methods

### Overview

The material and methods section is organized as follows. We first describe the new algorithm and compare it to the released version. This description focusses on the leaf area calculation and the tillering mechanism. Then, we describe the experiments and the data used to parametrize the genotypes and evaluate the model. The last three sections describe the procedure used to parametrize the genotypes, the criteria used for model evaluation, and the information about the software.

#### A. Leaf area calculation

The generic functions of the released pearl millet model are described in APSIM (2021). In the current studies, we enhanced the existing framework with two routines for different leaf area calculation (TPLA/ILA) connected with three different ways to reflect the tillering mechanism (fixed TPLA, fixed ILA, dynamic ILA). This allows the user to choose the level of model complexity based on the available data and based on the purpose of the simulation exercise. The pearl millet model new algorithm allows the user to choose between the TPLA and ILA approaches to model leaf area.

##### A.1. Total Plant Leaf Area (TPLA) approac

The total plant leaf area function is a simple way to describe the total plant leaf area at each point of crop growth and development. It can be expressed like that:

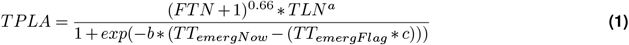

where *FTN* is the number of fertile tillers (need to be fixed by the user), *TLN* is the total final leaf number (with maximum leaf number set at 26 for both TPLA and ILA), *TT*_*emergNow*_ represents the cumulated thermal time from emergence to the actual time of the system, and *TT*_*emergFlag*_ is the cumulated thermal time from emergence to flag leaf.

The variables *a, b*, and *c* represent parameters that can take genotype specific values. Parameter *a* is the main stem coefficient (*pow. coeff. for TPLAmax*, table 3). Parameter *b* is the TPLA production coefficient that represents the breadth of the TPLA function (*curvature coeff for leaf area*, table 3). Parameter *c* is the TPLA inflection ratio that represents skewness of the TPLA function (*inflection coeff of TPLA curve*, table 3). In Figure 2 (a-c), we illustrate the TPLA function with the influence of the *a, b*, and *c* parameters on the leaf area. The TPLA approach is useful for users who want to parametrize the model without measuring the area of individual leaves. The TPLA function also allows to compare pearl millet with other crops models whose leaf area modelling is based on the same principle.

**Figure 2.**
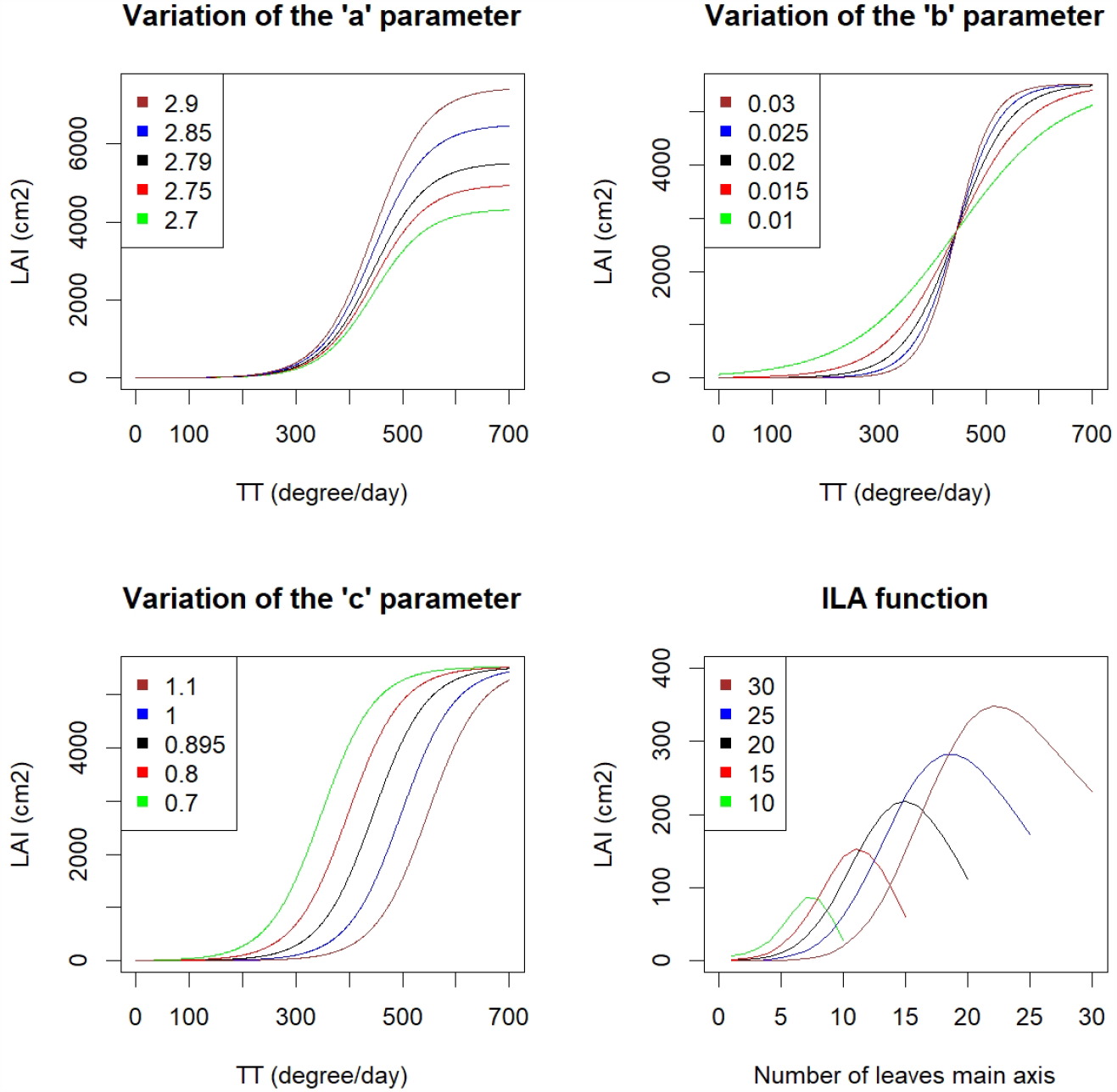
Illustration of the TPLA and ILA functions with the influence of the three genotype specific parameters and the total number of leaf (TLN): a) influence of parameter *a* representing the main stem coefficient on TPLA function, b) influence of parameter *b* representing the breadth of the function on TPLA function, c) influence of parameter *c* representing the skewness of the function on TPLA function; and d) ILA function for different values of TLN (10, 15, 20, 25, 30)

##### A.2. Individual Leaf Area (ILA) approach

The ILA approach reproduces the establishment of canopy based on individual leaf growth. It is a prerequisite to model tillers in a dynamic way (Van Oosterom et al., 2001a). The area of an individual leaf (*Y*) can be calculated using the following bell-shaped function for which parameters can be further expressed in terms of the total leaf number (TLN) of an axis (Dwyer and Stewart, 1986):

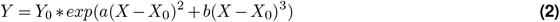

where *X* is the leaf ordinal position on the axis (1, 2, 3, …), *X*_0_ is the position of the largest leaf and *Y*_0_ the mature area of the largest leaf. *X*_0_ and *Y*_0_ can be further parametrized to introduce some genotype specificity in the leaf area calculation.

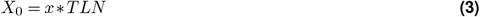

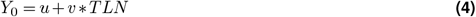

where *x* is a genotype specific parameter representing the *largest leaf multiplier, u* is the *intercept for largest leaf calculation*, and *v* is the *largest leaf area factor* (table 3). The released model also uses the ILA approach for leaf area. Compared to this model, the new algorithm includes possibility of a genotype specific parametrization of the largest leaf position calculation (parameter *x*). Figure 2 (d) contains an illustration of the ILA function for different value of TLN. The parameters *a* and *b* determine the breadth and the skewness of the bell-shaped curve, respectively. Given expressions and values from Birch et al. (1998), *a* and *b* were defined as:

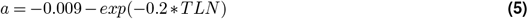

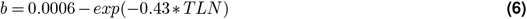

#### B. Tillering mechanism

In the released model, the default option is to model tiller appearance as a function of the thermal time. Tillering starts at 150 °*Cd* after crop emergence, at a rate of 34 °*Cd* per tiller. The number of emerged tillers is an input of the model and the maximum number of tillers is set to five. The rate of tiller appearance is not dependent on genotype. Tiller death is the result of insufficient resource capture to produce a panicle. A tiller will not produce any grain and die if its average growth rate between flag leaf appearance and grain filling is below 0.1 g/plant/day.

The extended tiller model gives some extra flexibility to the user by allowing a dynamic determination of the number of tillers, by relaxing their maximum number, and by allowing the use of genotype-specific tillering rate. It also revises the tillering algorithm by simulating tillering as a carbohydrate S/D ratio function based on actualized biological knowledge. The new tillering algorithm is based on the work of Kim et al. (2010a,b), using the simplified algorithm from Alam et al. (2014).

Tillers start to appear at 150 °*Cd* after emergence from the base of the third leaf when the fifth leaf (L5) is under development. The potential tiller number (*TTN*) is determined when L5 is fully expanded and is a function of the carbohydrate S/D ratio.

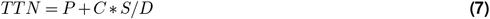

*P* is the expected propensity to tiller and *C* the propensity to tiller given extra unit of carbohydrate S/D (*propensity to tiller* and *tiller supply/demand slope*, table 3). The supply (S) is a product of: the average radiation per unit of thermal time over the period of L5 expansion, L5 area, and L5 phyllochron. D is calculated as the difference in area between L9 and L5. The basic principle builds on the estimation of carbon supply available for the main stem leaf growth (main carbohydrate demand) and the remaining carbohydrates could be allocated for tillers growth (Alam et al., 2014).

The tiller leaf size is directly related to the leaf size of the main stem with an empirically estimated reduction coefficient of 5% (*vertical offset for the largest leaf calc*., table 3). The first tiller largest leaf area is equal to 95% of the main stem, the second tiller to 90%, etc. Two mechanisms can cause tiller growth cessation. Tillers stop growing when the leaf area index (LAI) equals 0.65 (Lafarge et al., 2002) or when there is no more carbohydrate supply for tiller growth below a certain specific leaf area (SLA *g/m*^2^) threshold (carbohydrate are preferably allocated to the main stem). The SLA threshold is defined as *SLA* = *−*18.158*∗ LN* + 429.72 with values bounded between 150 and 400 *g/m*^2^.

The extended parametrization of the ILA function (*largest leaf multiplier, the largest leaf area factor*, and *intercept of the largest leaf calculation*) as well as the genotype specific tillering control are essential to build a more efficient dynamic tillering function of the extended pearl millet model. The released model already had a sort-of dynamic tillering algorithms implemented. However, this was done in a rather cumbersome way with different parametrisations for the plant culms with separate tillers grown as an inter-crop.

#### C. Data

The data were collected on two genotype cultivars (9444 and HHB 67-2) in four experiments with different plant densities. In the following sections, we called low density (LD), plant density equal to 6.7 plants/*m*^2^ and high density (HD), plant density equal to 12 or 13.3 plants/*m*^2^.

##### C.1 Genotype – 9444

9444 is the largest pearl millet selling hybrid produced by Bayer and was introduced in the 90s (Yadav and Rai, 2013). 9444 is characterized by a comparatively late duration (80-90 days) and lower tillering. 9444 was bread for better endowed environment and represents the crop type which is broadly cultivated in the B zone but is also well represented in the A and A1 zones because according to Asare-Marfo et al. (2010), it is among the 10 most popular varieties cultivated in Rajasthan.

##### C.2. Genotype -HHB 67-2

HHB 67 is one of the most adopted pearl millet hybrid released by Haryana Agricultural University in the 90s (Yadav and Rai, 2013; Nagaraj et al., 2012) and represents the crop type popular in drought-prone environments. HHB 67 is an extra-early variety (62-65 days) with high tillering propensity which makes it adapted to abiotic stress, and adapted to low rain, two characteristic conditions of the A1 zone. HHB 67-2 is an improved version of HHB 67 produced after the molecular introgression of resistance to downly mildew (Nagaraj et al., 2012).

##### C.3. Experiment details

The four experiments (Table 1 and 2) were conducted in 2015, 2016, and 2017 at Patancheru, India (17°45*′N*, 78°16*′E*). All experiments were sown with a row spacing of 60 cm and were irrigated once or twice every two weeks to avoid drought stress. In terms of phenotypic data (Table 2), we selected the following traits to evaluate the model prediction ability: flowering time was measured as the day when 50% of the plot reached the flowering stage, leaf area was measured as the total leaf area of four selected plants at different time points, biomass represented the total plant biomass of a one squared meter plot surface at different time points, 1000 grains weight as well as grain number where measured from the four selected plant at harvest, the total number of primary tillers (fertile and non-fertile) was measured at different time point on the four selected plants, grain yield was the total grain weight divided by the remaining plot surface at the end of the experiment.

**Table 1.** Experiments description.

| Experiments | Sow. date | Day length | Density | Genotypes |  |
| --- | --- | --- | --- | --- | --- |
| | | (h/day at emergence) | (plant/$m^2$) | 9444 | HHB 67-2 |
| 1-Pat LD 2015 | 04/02/2015 | 12.2 | 6.7 | x | x |
| 2-Pat HD 2015 | 04/02/2015 | 12.2 | 13.3 | x | x |
| 3-Pat LD 2016 | 08/02/2015 | 12.3 | 6.7 | x | x |
| 4-Pat HD 2017 | 22/02/2015 | 12.5 | 12 | x |  |

**Table 2.** Phenotypic traits measured during the experiments.

| Experiments | Traits |  |  |  |  |  |  |
| --- | --- | --- | --- | --- | --- | --- | --- |
|  | Flowering | Leaf area | Biomass | Gr. weight | Gr. Nb | tillers | Gr. yield |
| 1-Pat LD 2015 | x | x | x | x | x | x | x |
| 2-Pat HD 2015 | x | x | x | x | x | x | x |
| 3-Pat LD 2016 | x |  | x |  |  | x | x |
| 4-Pat HD 2017 | x |  | x |  |  | x | x |

**2015**: Experiment 1 and 2 were sown on the 4th of February 2015 with a density of 6.7 and 13.3 plants/*m*^2^, respectively. Three genotype replications were laid out as a split-plot design with density as the main factor. Concerning management, a basal fertilizer dose of 200 kg/ha single super phosphate and 50 kg/ha diammonium phosphate was applied. A top dressing of 100 kg/ha urea was applied three weeks after planting.

**2016**: Experiment 3 was sown on the 8th of February 2016 with a density of 6.7 plants/*m*^2^. The considered data were part of a larger experiment with 14 other genotypes planted in a randomized complete block design (RCBD) with two replications per genotypes. Concerning fertilization, a basal dose of 60 kg/ha single super phosphate and of 100 kg/ha calcium ammonium nitrate was applied.

**2017**: Experiment 4 was planted on the 22th of February 2017 with a density of 12 plants/*m*^2^. Those data were also part of a larger experiment laid out as an RCBD with five replications per genotypes. In this experiment only 9444 was observed. In terms of fertilization, a basal dose of 100 kg/ha of diammonium phosphate and 50 kg/ha of urea, and a top dressing of 50 kg/ha of urea were applied.

#### D. Genotype parametrization

The main genotype specific parameters were derived from the gathered data. It is generally advised to estimate the parameters in a gradual order of physiological complexity: phenology, canopy, tillering, and biomass partition.

##### D.1. Phenology

Next to the leaf area and tiller parameters, the pearl millet model also enables to determine genotype specific parameters controlling the plant phenology. For example, we estimated the parameter thermal time from the end of the juvenile phase to panicle initiation (*TTendjuvInit*, table 3) using the software Devel 2.0 (Holzworth and Hammer, 1996).

##### D.2. Canopy

For the TPLA method, the parameters *a, b*, and *c* from equation 1 can be empirically determined by optimization given observed values for *TPLA, FTN, TLN, TT*_*emergNow*_, and *TT*_*emergFlag*_. Concerning the ILA approach, the parameters *x, u*, and *v* can be empirically determined given observed leaf area and *TLN* values through an optimization of 2 after replacing *X*_0_ and *Y*_0_ by expressions 3 and 4, respectively.

##### D.3. Tillering

For the tillering function, the parameters *P* and *C* from equation 7 can be determined for each genotype by fitting a linear regression between the observed maximum number of tillers and the S/D curve. *P* and *C* are the regression intercept and slope.

##### D.4. Grain number and biomass parameters

Finally, two parameters associated with grain number and grain biomass: dry matter per seed (*dm*_*per*_*seed*) and maximum grain filling rate (*maxGFRate*, table 3) can be determined using empirical data and specific formulas. The *dm*_*per*_*seed* parameter is the ratio between growth rate between floral initiation and grain filling, defined as daily increase in biomass, and the total grain number.

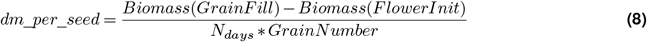

The *maxGFRate* parameter can be determined using the following formula:

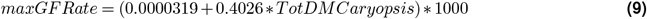

Where *T otDM Caryopsis* is the mass increase in [g/plant/grain/°*Cd*].

#### E. Model evaluation

To evaluate the new algorithm, we compared the observed and the predicted values using visualization techniques and metrics like the coefficient of determination (*R*^2^) and the d-index (Willmott-index), which accounts for data variability in both observed and predicted values (Wallach et al., 2018).

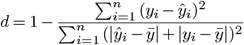

We first used scatter plots of the observed and predicted values which allowed us to check the alignment with the 1:1 line and observe if data fell into a 20% coverage interval around the 1:1 line. For development traits like the leaf area, the biomass, and the tillers we also made plot of observed and simulated values versus time. We compared prediction ability of different versions of the model (e.g. TPLA versus ILA version) using the root mean squared error (RMSE).

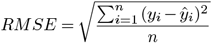

#### F. Software

For the moment, the presented new algorithm takes the form of add-on files available on request that can be integrated in the APSIM 7.10 environment.

## Results

### G. Genotype parametrization

In table 3, we listed the parameters of the two considered genotypes. The bold values represent the parameters that were determined using the observed data, which constitutes differences between those two plant types. The larger *TT endjuvInit* obtained for 9444 compared to HHB 67-2 (265 vs 175 °*Cd*) reflects the difference of crop duration, with HHB 67-2 being an extra-early variety and 9444 a longer variety. The grain filling parameters *dm*_*per*_*seed* and *maxGFRate* could be determined using data from experiment 2015 only. We could observe a small difference between the genotypes reflecting the smaller grains of HHB 67-2 compared to the one of 9444.

The ILA parameters were also determined using only data from experiment 2015. According to those values, the largest leaf of 9444 tends to be positioned at lower position compared to HHB 67-2. The area of the largest leaf (*Y*_0_) of 9444 decreases when TLN increases while for HHB 67-2 it increases. Finally, concerning the tillering parameter, we could notice that the expected larger propensity to tiller of HHB 67-2 compared to 9444 was reflected in those coefficients (PTT = 2.29 (9444) and 3.4 (HHB 67-2)).

### H. New algorithm evaluation

In figures 3 and 4, we plotted the observed against predicted values of flowering time, leaf area, number of tillers, biomass, and yield for the two genotypes. Table 4 first four columns contain the *R*^2^ and *d* values obtained for the evaluation of all traits.

**Table 3.**
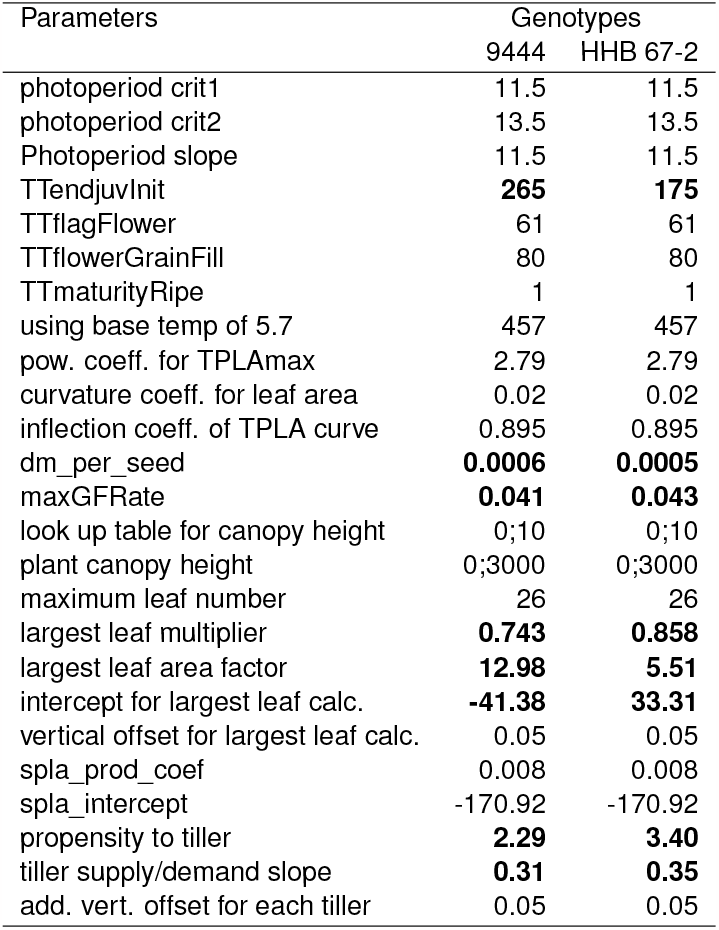
Genotype parameters of the APSIM pearl millet model new algorithm. Parameters that are different between two genotypes (bolded)

| Parameters | Genotypes |  |
| --- | --- | --- |
|  | 9444 | HHB 67-2 |
| photoperiod crit1 | 11.5 | 11.5 |
| photoperiod crit2 | 13.5 | 13.5 |
| Photoperiod slope | 11.5 | 11.5 |
| TTendjuvInit | **265** | **175** |
| TTflagFlower | 61 | 61 |
| TTflowerGrainFill | 80 | 80 |
| TTmaturityRipe | 1 | 1 |
| using base temp of 5.7 | 457 | 457 |
| pow. coeff. for TPLAmax | 2.79 | 2.79 |
| curvature coeff. for leaf area | 0.02 | 0.02 |
| inflection coeff. of TPLA curve | 0.895 | 0.895 |
| dm_per_seed | **0.0006** | **0.0005** |
| maxGFRate | **0.041** | **0.043** |
| look up table for canopy height | 0;10 | 0;10 |
| plant canopy height | 0;3000 | 0;3000 |
| maximum leaf number | 26 | 26 |
| largest leaf multiplier | **0.743** | **0.858** |
| largest leaf area factor | **12.98** | **5.51** |
| intercept for largest leaf calc. | **-41.38** | **33.31** |
| vertical offset for largest leaf calc. | 0.05 | 0.05 |
| spla_prod_coef | 0.008 | 0.008 |
| spla_intercept | -170.92 | -170.92 |
| propensity to tiller | **2.29** | **3.40** |
| tiller supply/demand slope | **0.31** | **0.35** |
| add. vert. offset for each tiller | 0.05 | 0.05 |

**Table 4.**
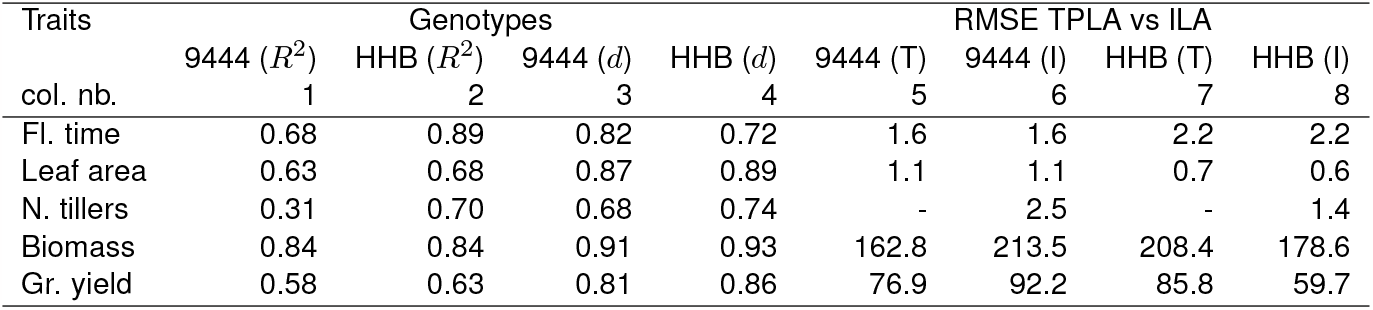
Extended model validation results: *R*^2^ for the regression of the observations versus model predictions, *d* values, RMSE for the comparison between TPLA and ILA.

**Figure 3.**
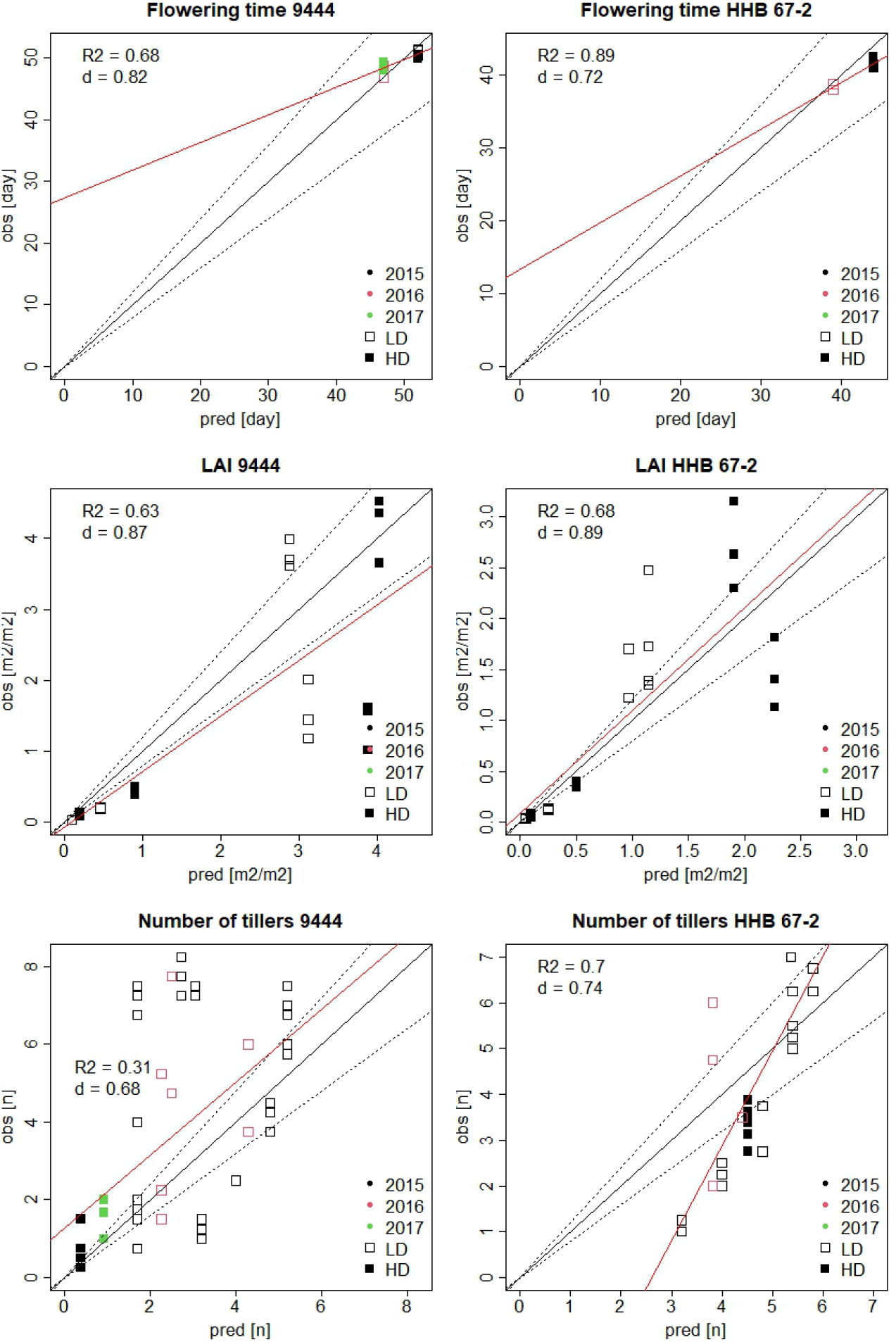
Scatter plot of observed versus predicted flowering time, leaf area, and number of tillers

**Figure 4.**
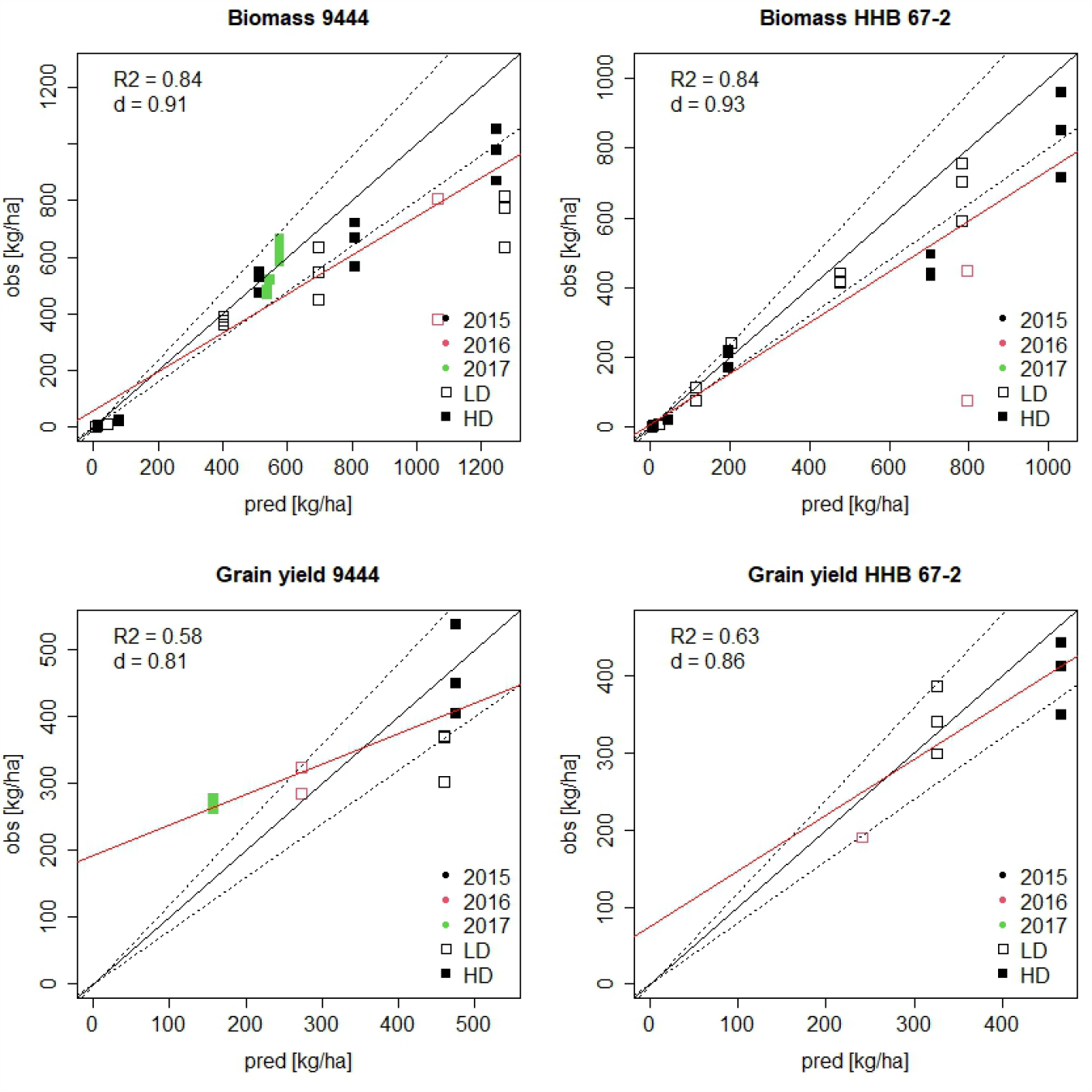
Scatter plot of observed versus predicted grain yield and flowering time

Looking at flowering time, biomass, and grain yield on figures 3 and 4, we noticed that most of the points are close to the 1:1 line or fall within the 20% coverage interval. The results for 9444 grain yield are slightly less good with the prediction of year 2017 being underestimated and the one of 2015 LD being overestimated. Those results are reflected in the *R*^2^ values with a reduced yield prediction ability (*R*^2^ = 0.58 (9444) *and* 0.63 (*HHB* 67 *−* 2)). However, the *d* values obtained for grain yield that took into account the variability in observed data were better (0.81-0.86).

Concerning the leaf area and the number of tillers, we could observe more points falling outside the 20% coverage interval. For example, concerning 9444 number of tillers, we noticed several large observed values that were not well predicted by the model (upper left side). This divergence was reflected in the low *R*^2^ value (0.31).

Figures 5 and 6 represent plot of the observed and predicted values over time per experiment of leaf area, biomass, and number of tillers for genotype 9444 and HHB 67-2, respectively. We could generally consider that the dynamic of those traits was reasonably well captured by the model with *d* values between 0.68 and 0.93. The dynamic of leaf area development is fairly described by the model, but for both genotypes we could observed some important differences between the observed and predicted leaf area at later stages. For example, for 9444, after reaching its maximum, the observed leaf area dropped more quickly than the prediction. Globally, those discrepancies explain that for leaf area, the *R*^2^ values are good but not extremely high (0.63 and 0.68).

**Figure 5.**
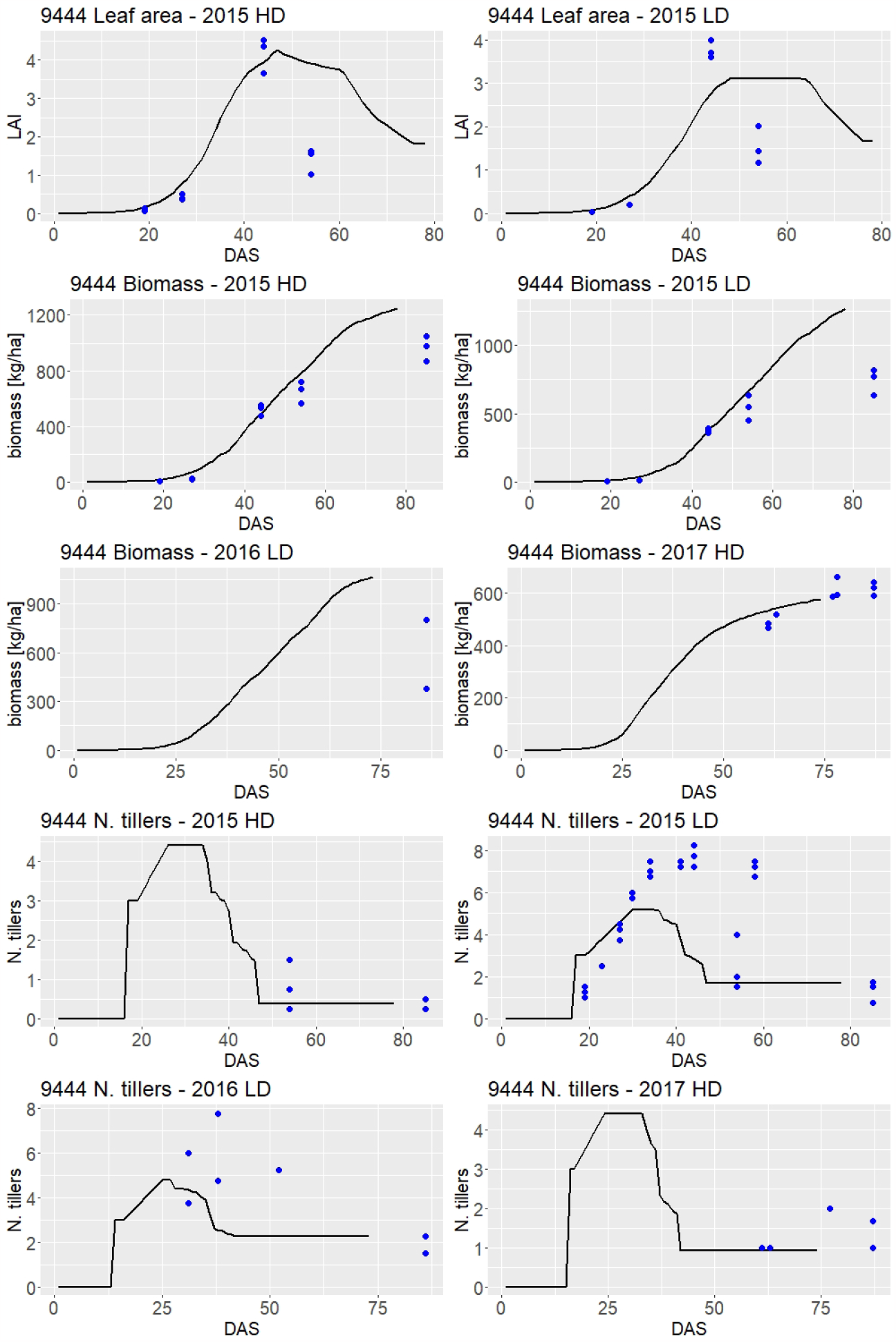
Plot of 9444 observed and predicted leaf area, biomass, and number of tillers values over time

**Figure 6.**
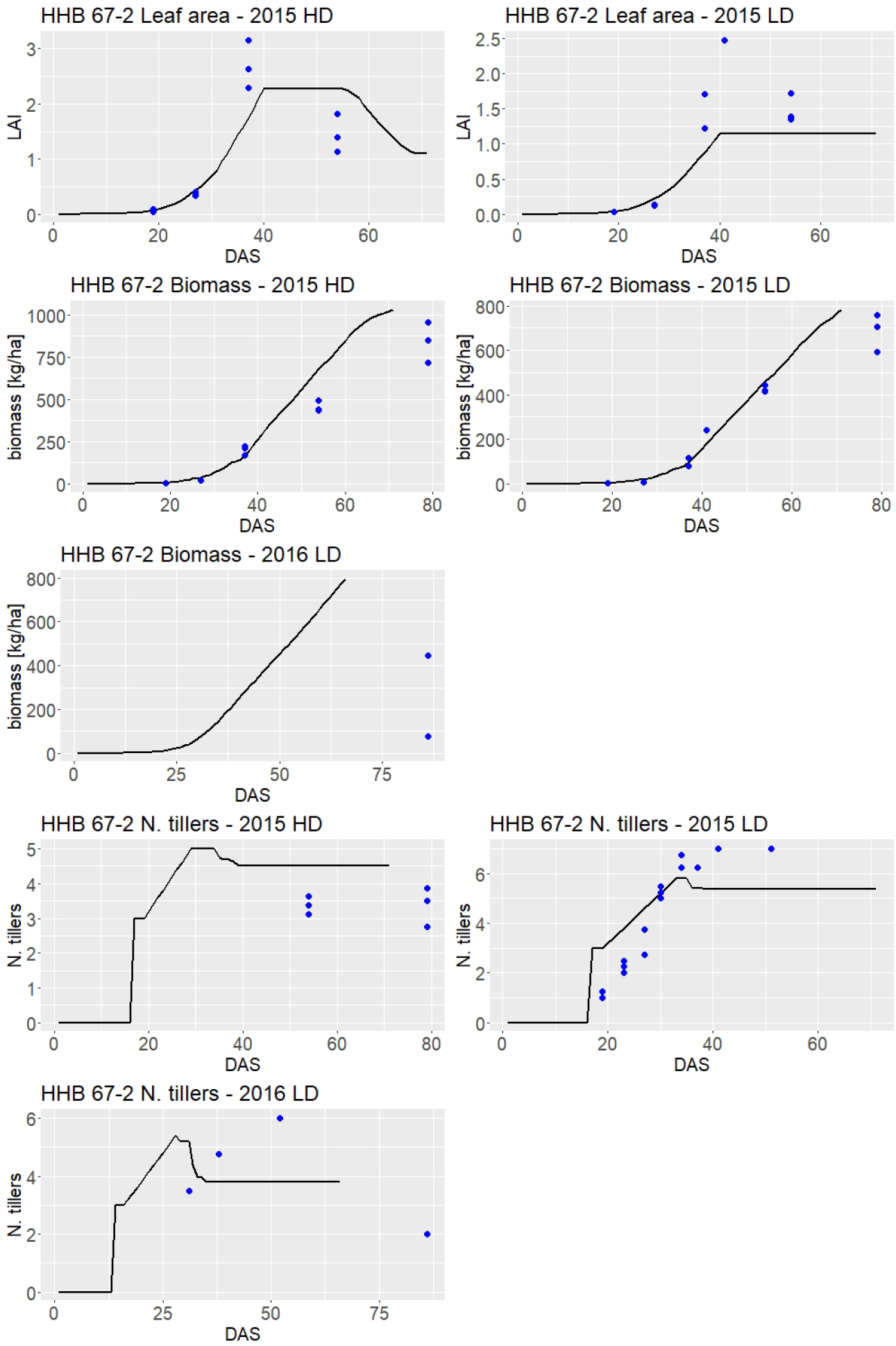
Plot of HHB 67-2 observed and predicted leaf area, biomass, and number of tillers values over time

Concerning biomass, we noticed on figures 5 and 6 that the model captured well the biomass accumulation process. Some divergence between the predictions and the observations could be observed concerning the biomass at harvest. In several cases (e.g. 9444 2015 and HHB 67-2 2016 LD), we could observe a systematic overestimation of the biomass by the model.

Looking at the number of tillers, we could see that the model captured well the initial growth phase (e.g. 9444 and HHB 67-2 2015 LD) but tended to decline faster than what was observed. However, the final number of tillers was relatively well predicted by the model for 9444 but was slightly overestimated for HHB 67-2.

### I. Comparison TPLA vs ILA approach

For each trait, we calculated the RMSE to compare the prediction ability of the extended model using the ILA and the TPLA functions. The columns five and six of table 4 describe the TPLA versus ILA difference for 9444 while the column seven and eight the differences for HHB 67-2. According to those results, leaf area prediction is similar between those two options. We could observe some significant difference between ILA and TPLA for biomass and yield estimation. However, we could not determine if one option performed generally better than the other because for 9444, TPLA obtained smaller difference for biomass and yield prediction, while for HHB 67-2, it was the opposite, ILA performed better than TPLA. By definition, TPLA approach can not predict the number of tillers.

## Discussion

The main goal of this study was to present and evaluate a new algorithm for pearl millet modelling in APSIM. Based on the presented results we can say that globally the new algorithm describes reasonably well the observed data but that some discrepancies emphasize the need to continue the development of pearl millet crop modelling.

### J. The challenge of tillering cessation modelling

According to the results, the most important challenge is the modelling of tiller cessation. For HHB 67-2, the model could predict tiller growth and capture the persistence of a large number of tillers. For genotype 9444, we could observe an important tillering growth before a reduction. The transition between a large number of tillers and the remaining one happened faster in the model, which explain some larger differences, especially in experiment 2015 LD for 9444. Those large tiller numbers data represent an important part of the divergence between the predictions and the observations (figure 3 9444 number of tillers upper left). Indeed, if we remove the observations above 6.5 tillers in experiment 2015 LD for 9444 (13 values out of 50), the *R*^2^ for tiller would increase from 0.31 to 0.61. In that case, 9444 tillering prediction would be equivalent to the one of HHB 67-2.

A too liberal count of the number of tillers could explain the lower prediction for 9444. The fact that we observed tiller number above six for 9444, which is supposed to be a low tillering variety support that hypothesis. A too liberal count of tillers specific to LD experiment could be explain by the fact that in LD situations, tillering become more profuse (Van Oosterom et al., 2001b). Such a large number of tillers could make it more difficult to distinguish between primary and secondary tillers. Since the model only predicts primary tillers, a part of the divergence could be explained by secondary tillers wrongly considered as primary tillers. At the end, the final number of tillers of 9444 was predicted with a good accuracy. Therefore, for this genotype the difficulty is concentrated in the dynamic of tiller cessation, which is an important challenge, especially in a species like pearl millet with a larger propensity to tiller compared to crops like sorghum.

We should also emphasize that the tillering algorithm used in this study come from work on sorghum Kim et al. (2010a); Alam et al. (2014). The adaptation of such framework to pearl millet, which is characterized by a larger tillering propensity could required more specific work. For example, the determination of genotype tiller propensity and relationship between number of tillers and carbohydrate supply could be improved by running experiment using a larger amount of genetic and environmental variability. To the extend of our knowledge, this work is a first attempt since pearl tillering initial work by Van Oosterom et al. (2001a). Despite some evident progresses like the possibility to predict tillers in a dynamic way, pearl millet tillering still deserve some extra attention.

Further parametrization work on specific leaf area (SLA) could be useful to improve tiller cessation mechanism because tiller cessation is related to the impossibility for leaves to reach the minimum thickness, which is related to maximum SLA. Since this parameter was not specifically covered in this study, extra work on that direction could represent the future steps.

### K. Other limitations

Another point to emphasize is the almost systematic overestimation of the biomass at harvest by the model. Such a systematic result could be due to systematic underestimation of the observed biomass at harvest in the phenotyping procedure. Finally, we also would like to emphasize that the experiments were mainly run at the same time, which limits the possibility to cover an appropriate photoperiod range. Similarly, compared to other studies (e.g. Van Oosterom et al. (2001a)), the covered plant density is reasonable but it could be extended. Some extra data collection at different photoperiod time with extended plant density range would also help to strengthen the results observed in this study.

### L. Conclusions

To conclude, we consider that, despite some margins of improvement, the presented results represent valid steps for the improvement of pearl millet modelling. Our results validate the modifications done on the released APSIM pearl millet model to improve the existing leaf area and dynamic tillering modelling. The new algorithm allowing the definition of new genotype specific parameters gave us the opportunity to parametrize two important genotypes for Indian pearl millet production. Therefore, we consider that, in parallel of extra validation work, the extended model could be tested for larger scale predictions using other station data and/or some district data with good confidence in weather and soil data input. The possibility to apply the general ILA concept to pearl millet make it comparable to other crop algorithm like sorghum or maize. This is an important step to realize fair comparison between those crops within the same simulation environment.

### M. Funding

This work was supported by a grant from the Swiss National Science Foundation (Postoc.Mobility grant no: P500PB_203030). Dr Kholova contribution was financed by an internal grant agency of the Faculty of Economics and Management from the Czech University of Life Sciences Prague (Grant Life Sciences 4.0 Plus no. 2022B0006)

## Notes

### Competing Interest Statement

The authors have declared no competing interest.

### Summary of Updates

Addition of a section about funding.

